# Sleep-wake states are encoded across emotion-regulation regions of the mouse brain

**DOI:** 10.1101/2024.09.15.613104

**Authors:** Kathryn K. Walder-Christensen, Jack Goffinet, Alexandra L. Bey, Reah Syed, Jacob Benton, Stephen D. Mague, Elise Adamson, Sophia Vera, Hannah Soliman, Sujay Kansagra, David Carlson, Kafui Dzirasa

## Abstract

Emotional dysregulation is highly comorbid with sleep disturbances. Sleep is comprised of unique physiological states that are reflected by conserved brain oscillations. Though the role of these state-dependent oscillations in cognitive function has been well established, less is known regarding the nature of state-dependent oscillations across brain regions that strongly contribute to emotional function. To characterize these dynamics, we recorded local field potentials simultaneously from multiple cortical and subcortical regions implicated in sleep and emotion-regulation and characterize widespread patterns of spectral power and synchrony between brain regions during sleep/wake states. First, we showed that these brain regions encode sleep state, albeit to various degrees of accuracy. We then identified network-based classifiers of sleep based on the combination of features from all recorded brain regions. Spectral power and synchrony from brain networks allowed for automatic, accurate and rapid discrimination of wake, non-REM sleep (NREM) and rapid eye movement (REM) sleep. When we examined the impact of commonly prescribed sleep promoting medications on neural dynamics across these regions, we found disparate alterations to both cortical and subcortical activity across all three states. Finally, a we found that a stress manipulation that disrupts circadian rhythm produced increased sleep fragmentation without altering the underlying average brain dynamics across sleep-wake states. Thus, we characterized state dependent brain dynamics across regions canonically associated with emotions.

**Significance Statement:** Sleep and emotion regulation are known to be intertwined at the level of behavior and in neuropsychiatric illnesses. Here, we examined how brain regions involved in emotion regulation encode wake and sleep states by performing multi-site electrophysiological recordings in mice. We developed sleep-wake state classifiers that rapidly labeled sleep-wake states from brain activity alone. We then identified how commonly prescribed sleep-inducing medications have unique impacts on brain activity throughout these emotion-regulation regions. Finally, we explored the impact of circadian rhythm disruption on sleep architecture and brain activity. Together, these data shed light on how brain regions which regulate emotion behave during sleep so that one day, treatments to improve both sleep and emotional well-being may be developed.

## Introduction

Sleep is composed of fundamental states with distinctive behavioral, physiological, and neurophysiological signatures that are conserved across mammalian species (Lo et al., 2004; Keene and Duboue, 2018). Wake is a state of behavioral arousal with low-amplitude mixed frequency brain activity and high muscle tone. Sleep is a reversible state of behavioral quiescence, which can be further decomposed into rapid eye movement (REM) sleep, defined by low-amplitude, mixed mixed-frequency oscillations with increased theta and gamma brain activity and absence of muscle tone, and non-REM (NREM) sleep, defined by high-amplitude delta frequency brain activity and low muscle tone (Aserinsky and Kleitman, 1953; Kohn et al., 1974; Brown et al., 2012). Inadequate timing, quality, or quantity of sleep affects many aspects of health and disease, from cognition (Killgore, 2010) to immune function (Besedovsky et al., 2019). Impairment of sleep is a common and debilitating symptom shared across numerous neuropsychiatric disorders, especially in disorders of emotion regulation (Baglioni et al., 2016). Overlap in the regulation of sleep and mental well-being has been identified from the level of genes (e.g. genome-wide association studies linking molecular clock genes to bipolar and major depressive disorders; (Jagannath et al., 2017) to population-level epidemiology (e.g. higher rates of depression in night-shift workers (Jagannath et al., 2017). In experimental models, manipulating sleep quantity or quality can alter emotion regulation (Goldstein and Walker, 2014). There is evidence to suggest that sleep and emotion regulation overlap also occurs at the level of brain networks (Mu and Huang, 2019), though limited studies examine how emotion-regulation brain networks behave during sleep states or how sleep-altering treatments impact these brain dynamics.

There has been significant progress, largely in rodent models, in understanding how different populations of neurons distributed throughout the brain exert widespread effects to regulate transitions into, or maintenance of, sleep wake states, including forebrain cholinergic neurons, hypothalamic histaminergic and orexinergic neurons, and brainstem serotonergic, dopaminergic and noradrenergic neurons (Lu et al., 2006; Brown et al., 2012; Mahoney et al., 2019). These distributed ensembles of neurons work together to organize sleep architecture. Indeed, several studies using parallel recordings from multiple brain regions have demonstrated the utility of studying the regulation of sleep across distributed brain regions (Gervasoni et al., 2004; Takahashi et al., 2009; Zhang et al., 2022). Recent work using single- and multi-region electrophysiological data in mice demonstrated the ability to decode sleep-wake states using only milliseconds of neural activity (Parks et al., 2024). Though this study was largely limited to decoding patterns within (but not across) subjects, these findings indicated the robust signatures of sleep across the brain (Parks et al., 2024).

Similarly, the brain regions implicated in emotion and mood regulation are numerous and widely distributed, including cortical, limbic, striatal, midbrain, and brainstem areas (Nestler et al., 2002; Hultman et al., 2016; Hultman et al., 2018). There is notable overlap in various neuromodulatory systems regulating emotion and sleep, including cholinergic neurons, dopaminergic neurons, serotonergic neurons, and noradrenergic neurons, which may indicate shared neuronal networks underlying regulation of emotions and sleep-wake biology (Dzirasa et al., 2006; Brown et al., 2012; Mu and Huang, 2019). Here, we focus on a set of eight regions across the brain that are conserved between rodents and humans, implicated in emotional regulation, and are reliably accessible using standard electrophysiological microwires (Mague et al., 2022; Hughes et al., 2024). Specifically, we record from the cingulate cortex (CxCg), prelimbic cortex (CxPrL), infralimbic cortex (CxIL), nucleus accumbens (core and shell grouped together; NAc), amygdala (basolateral and central grouped together; Amy), medial dorsal thalamus (ThalMD), ventral hippocampus (HippV), and ventral tegmental area (VTA). All these regions, either in isolation or as components of neural circuits, have also been implicated in sleep biology. For example, basolateral Amy is critical for the transition between NREM and REM (Hasegawa et al 2022) and lesions of the central nucleus of the Amy decreased REM (Tang et al 2005). CxCg is associated with awakening in insomnia (Guo et al 2023, Li et al 2024), lesions in the prefrontal cortex including PrL and IL are associated with decreased REM sleep and increased sleep fragmentation (Chang et al 2014), and pyramidal neurons in IL have their highest activity during REM (Hong et al 2024). The core region of NAc can be activated to induce NREM sleep (Oishi et al 2017) and lesioning of NAc core and shell increases wakefulness (Qiu et al 2012). Lesion of ThalMD increased wakefulness in cats (Marini et al 1988) but decreased wakefulness in rats (Mallick et al 2020). HippV plays a critical role in emotional memory consolidation during sleep (Pronier et al 2023). Finally, GABAergic neurons in VTA promote NREM sleep (Chowdhury et al 2019). Though these findings clearly establish the role of these individual brain regions in sleep, it remains unclear how these regions are modulated in concert across sleep-wake states. Addressing such a question requires studies examining neural activity at the seconds-to–minutes timescale, which comprises the typical duration of a sleep state for mice and humans (Arrigoni and Saper, 2014). Such studies have been lacking as prior work has largely focused on sleep-related molecular and cellular activity on the millisecond-to–second timescale or circadian rhythms of sleep-wake behavior on the hours-to–days timescale.

Here we begin to close this gap by characterizing the brain oscillatory patterns that are reliably observed within and across emotional regions during sleep-wake states. We achieved this by applying supervised machine learning techniques to local field potentials (LFPs) recorded simultaneously from multiple brain regions in mice (Hultman et al 2018, Mague et al 2022, (Walder-Christensen et al., 2024). This approach yields electrical functional connectome-based (electome) networks, which integrate widespread neural activity from the milliseconds-to-minutes timescale. The activity of an electome network, defined by the co-expression of its component circuits across seconds of time, can then be used for brain-state classification, Thus, this approach enables brain states to be rapidly determined from long-term recordings (Talbot et al., 2023). We trained our electome-based classifier based on power and synchrony (cross-power) in 8 cortical and subcortical brain regions, and we validated it against EMG and brain-based measures traditionally used for distinguishing sleep-wake states. Finally, we characterized the impact of pharmacological manipulations, commonly prescribed for emotional dysregulation and sleep, on regional brain dynamics, and we investigate the impact of chronic stress on sleep brain dynamics using the EMG-free sleep-wake classifier.

## Methods

### Mice and husbandry

All care and experimental procedures were conducted according to IACUC-approved protocols at Duke University. Male and female C57BL/6J (C57; JAX strain s000664) mice purchased from the Jackson Labs (Bar Harbor, ME) were used to train (n=9 male) and validate (n=5 male [mice utilized again for circadian stress]/4 female) the model and for psychopharmacology (n=5 male mice) and circadian stress manipulation experiments (n=9 male mice). Mice were maintained in single sex 12-hour light: 12-hour dark cycle (except for animals undergoing chronic light stress, see below) with food and water *ad libitum*. Mice were group- housed (<=5 mice per cage) except during habituation and sleep recordings (see below). Studies were conducted with mice aged 9 to 30 weeks, and mice were recorded in a pseudo-home cage (see below for details). Animals were excluded if electrode tracts were identified outside of the targeted regions; 4 mice excluded based on this criterion from pharmacological manipulation study and are not included in the n reported for the analysis. Mice requiring EMG were excluded if EMG power signal was visibly saturated across all frequencies. Five mice excluded based on this criterion: 1 from the training dataset and 4 from the chronic stress implanted dataset, both of which are not included in the n reported for the analysis. Three mice did not survive the surgery or post-operative period and were thus not included in the n reported for the analysis.

### Electrode implantation surgery

39 mice (ages 8 weeks to 12 weeks) were anesthetized with 1% isoflurane, placed in a stereotaxic device, and metal ground screws were secured above the cerebellum and anterior cranium. The recording bundles designed to target basolateral and central amygdala (Amy), medial dorsal thalamus (ThalMD), nucleus accumbens core and shell (NAc), ventral tegmental area (VTA), medial prefrontal cortex (mPFC), and ventral hippocampus (HippV) were centered based on stereotaxic coordinates measured from bregma (Amy: -1.4mm AP, 2.9 mm ML, -3.85 mm DV from the dura; ThalMD: -1.58mm AP, 0.3 mm ML, -2.88 mm DV from the dura; VTA: -3.5mm AP, ±0.25 mm ML, -4.25 mm DV from the dura; HippV: -3.3mm AP, 3.0mm ML, -3.75mm DV from the dura; mPFC: 1.62mm AP, ±0.25mm ML, 2.25mm DV from the dura; NAc: 1.3mm AP, 2.25mm ML, -4.1 mm DV from the dura, implanted at an angle of 22.1°). To target cingulate, prelimbic, and infralimbic cortex, a mPFC bundle was assembled by building a 0.5mm and 1.1mm DV stagger into the electrode bundle microwires. NAc core and shell were targeted in a single bundle with a 0.6mm DV stagger. Basolateral and central amygdala were targeted by building in a 0.5mm ML + 0.25mm DV stagger into the Amy bundle. Animals were implanted bilaterally in mPFC and VTA. All other bundles were implanted in the left hemisphere. A 20um wire was inserted into the trapezius muscle to record muscle activity for a subset of mice used to acquire ground truth sleep labels along with multi-site electrophysiology. A subset of the mice had additional brain regions recorded that were not used in these studies. Five of the male mice used to generate the training dataset had wires placed in dorsal hippocampus. All of the chronic circadian stress mice (which were also used as out of sample validation mice) had EMG and wires targeting anterior insular cortex, dorsal hippocampus, orbital frontal cortex, dorsal lateral striatum, dorsal medial striatum, lateral habenula, motor cortex, sensory cortex, substantia nigra pars reticulata, and visual cortex. To mitigate pain and inflammation related to the procedure, all animals received carprofen (5 mg/kg, s.c.) injections once prior to surgery and then daily for three days following surgery.

### Electrode placement confirmation

Histological analysis of implantation sites was performed at the conclusion of experiments to confirm recording sites used for neurophysiological analysis. Animals were perfused with 4% paraformaldehyde (PFA) and brains were harvested and stored for 24 hrs in PFA. Brains were sliced at 40μm using a vibratome (Leica) and stained using DAPI (ab104139, Abcam, Cambridge, MA) in the mounting media. Images were obtained using an Olympus Slide Scanner (VS200) at 4x.

### Neurophysiological data acquisition

Similar to our prior work (Mague et al., 2022), mice were connected to either an M or mu headstage (Blackrock Microsystems, UT, USA) and placed into a cage with food and water.

Mice were habituated to single housing for 2 days prior to recording. Mice were given extra enrichment including a chew bone, a wooden block, and a cup to hold food pellets.

On the day of recording, mice were placed into new cages with a small amount of bedding from their previous cage and allowed to acclimate for 3–6 hours. Subsequently, neural activity was sampled at 30kHz using the Cerebus acquisition system (Blackrock Microsystems Inc., UT) for at least 24 hours inclusive of a full light/dark cycle. Local field potentials (LFPs) were bandpass filtered at 0.5–250Hz and stored at 1000Hz. An online noise cancellation algorithm was applied to reduce 60Hz artifact. Only the final 24 hours of recording were utilized in the subsequent analyses. Neurophysiological recordings were referenced to a ground wire connected to ground screws in the skull.

### Sleep/Mood Medication Exposure

After surgical recovery of at least 2 weeks, implanted mice were singly housed for 2 days prior to recording. Mice were recorded for 24 hours following acute administration of a medication via i.p. injection (diazepam 3mg/kg or trazodone 40mg/kg solubilized in 10% DMSO + 0.9% saline). Mice were given a one-week washout period before administration of the other medication or saline control (containing 10% DMSO). Power and synchrony features during the initial 2 hours were averaged for each sleep-wake state based on network-classification labels.

### Chronic Stress Induction by Aberrant Light:Dark Cycle

Mice were single housed and connected to the electrophysiology recording system. The 24 hours prior to any change in their light cycle is considered the baseline recording (12:12 light-dark [T24]). Then, the mice were exposed to chronic aberrant light interruption using a 3.5:3.5 light dark (T7) cycle for 14 days (LeGates et al., 2012). After 14 days, the mice were immediately returned to the T24 light condition and the following 24 hours were analyzed for the post-stress session.

### LFP Preprocessing and Feature Generation

Power spectral density plots were calculated for each contact and inspected manually for the removal of poor channels (i.e., saturated). Outlying samples from each channel were identified automatically as those points in a 30Hz-highpassed signal with an absolute value above 20 median absolute deviations of the channel’s activity. Individual channels in the same region of interest were then averaged in the time domain (NAc core and shell were averaged together and basolateral and central amygdala were averaged together). Then the LFPs were segmented into 2-second windows and cross-power spectral densities were calculated using Welch’s method independently for each window, both within regions (power) and between regions (cross-power, or synchrony). 29 frequency bins linearly spaced between 0 and 54.7 Hz are collected. Power and synchrony features were normalized by their median value over all frequencies and regions, within each mouse for each recording.

### EMG-informed Cluster-based Classification

A graphical interface was developed to annotate 2-second windows into wake, REM, and NREM sleep bouts by two-dimensional state mapping of theta, gamma, and delta power from single-channel LFPs with EMG atony used to delineate overlapping REM versus wake recordings based on previously defined electrophysiological properties (Gervasoni et al., 2004). EMG power was calculated as the sum of the signal’s average power per frequency bin in the 30- 60Hz band and in the 60-250Hz band and was plotted on the y-axis (Dzirasa et al., 2006). Root mean square of LFP power was based on a single selected electrode site – here we use prelimbic cortex, ventral hippocampus, or dorsal hippocampus – and was plotted on the x-axis. The wake cluster was identified based on high EMG power, the NREM cluster was based on low EMG but high LFP power, and the REM cluster was based on low EMG power and low LFP power.

Manual discrimination of clusters was done by drawing a polygon around the cluster and assigning it to a particular state, after which a user manually circles the three regions corresponding to wake, NREM, REM, and unknown sleep-wake states. A second plot based on spectral LFP power ratios (x-axis: (0.5-4.5Hz)/(0.5-9Hz); y-axis: (0.5-20Hz)/(0.5-55Hz) was plotted. Clustering was refined using the second plot where the upper cluster (high power in <4.5 Hz frequencies) was defined as NREM. We observed high mixing of wake and REM in the bottom cluster and thus it was not used to distinguish between those states. All points that were outside of the hand-drawn polygons were considered unlabeled and were not used to calculate the accuracy of the EMG-free classifiers. Clusters were identified for data in 60-minute segments to allow for optimal balance of labeling efficiency and discriminability between clusters.

### Hand-Labeling of Sleep Labels

The EMG-informed cluster labels were validated against expert hand-labeling of sleep-wake states for a subset of recordings utilizing LFP traces from prelimbic cortex, ventral hippocampus, and the EMG. Individual 30- or 60-minute recordings were inspected and annotated with sleep-wake state labels based on heuristics established in the literature (Silber et al., 2007): low-amplitude, mixed-frequency brain oscillations along with increased muscle power in wake; high-amplitude, delta-predominant activity in cortex and hippocampus with low muscle power in NREM sleep; and low-amplitude, mixed-frequency oscillations with increased theta and gamma power in cortex and hippocampus with very low muscle power in REM sleep epochs, which were preceded by NREM epochs. Hand labels were annotated with a granularity of 2s to match the automated sleep classifier and was chosen to optimally minimize multi-labeled windows. Example labels of each state, transitions, and difficult-to-classify states were discussed with a sleep medicine specialist prior to establishing scoring rules. There was no coordination between the labelers after establishing the scoring rules.

### Inter-rater Agreement

Three raters classified sleep-wake states for each 2s window in 4 1-hour recordings of mouse LFPs and EMG. For this evaluation, there was no option to leave a window unclassified.

The inter-rater agreement was quantified by calculating Cohen’s kappa for each pair of label sets using scikit-learn’s cohen_kappa_score function.

### Supervised Autoencoder Sleep-Wake State Classifier

A supervised autoencoder (SAE) model was trained to predict EMG-informed cluster-based sleep labels (wake vs. NREM vs. REM) in 2s windows. This follows the logic and methods described by Talbot et al. 2023. The SAE converts the observed neural data to a low-dimensional feature representation that both accurately summarizes the neural data and is predictive of sleep-wake state. The parameters are learned to both predict well and reconstruct the observed neural features. More specifically, the SAE minimizes the following objective:

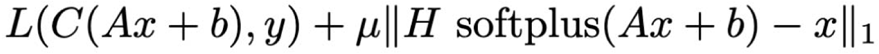

where is the vectorized neural features (power and cross-power) associated with a single window, and the matrix and vector define a linear transformation from the high-dimensional neural feature space (d=1856) to a low-dimensional summary feature representation (d=32), termed “electome scores.” The first term in the objective is the prediction loss, where denotes cross-entropy loss with logits a and sleep-wake state label vector and is a diagonal matrix with 3 rows, one for each sleep-wake state. The second term in the objective is the reconstruction loss, for which the summary features (electome scores) are rectified with a softplus nonlinearity and subsequently projected to the high-dimensional neural feature space by the nonnegative matrix, defining the generative model associated with nonnegative matrix factorization. μ > 0 determines the importance of the reconstruction relative to sleep label prediction. Each row of H is a nonnegative vectorized rank-1 tensor (regions-by-regions-by- frequencies) that associates each electome score with a factor, termed an “electome factor,” in the high-dimensional neural feature space. The first 3 rows of H are termed the supervised electome factors because they correspond to each of the three sleep-wake state logits and can therefore be used to interpret the neural features that discriminate between the sleep-wake states, whereas the remaining rows are unsupervised electome factors. It is common for SAEs to use complex mappings defined by multilayer neural networks to map neural features to electome scores (Talbot et al 2023), but here we use a simple linear mapping with a nonlinearity because we found this to be sufficient to obtain good performance.

To describe the classification performance of the SAE, we report balanced accuracy, which is the average recall of the classifier across all three sleep-wake states. This metric weights the classification of each of the three sleep-wake states equally, not in proportion to the portion of time spent in each state.

We used leave-one-mouse-out cross-validation to perform model selection and estimate model generalization on 9 training mice to ensure rigorous generalization estimates (Krstajic et al., 2014). We used three randomly shuffled splits of 6 training mice and 2 validation mice and considered hyperparameter settings of μ=10^-1, 10^-2, 10^-3 for the interior loop of the nested cross-validation. After obtaining an unbiased estimate of performance and selecting the hyperparameters, we retrained on all 9 mice with the chosen setting. This is the only model considered subsequently (and had no access to the rest of the validation studies during training). The same nested cross-validation procedure was used when considering the single-region models.

### Sleep Architecture Analyses

To integrate temporal information into the SAE’s per-window label prediction, a top-*k* Viterbi algorithm was applied to the label sequences. This method takes as input a transition matrix that describes the probability of transitioning from a given state to every other state in the next window (Viterbi, 1967). This matrix was estimated using several hours of hand-labeled sleep-wake states and collecting the empirical state transitions from each 2-second window to the next. The top-*k* Viterbi algorithm then gives the k most likely state sequences, given the SAE- produced per-window state probabilities and expected transition statistics. Sleep bout and transition statistics were calculated using these “smoothed” label sequences with *k*=10.

### Summarizing LFP Features That Distinguish Experimental Conditions

To compare LFPs within the same sleep-wake state, but across various experimental conditions (male vs. female, drug vs. saline, and before vs. after chronic sleep deprivation), we test for statistical differences of LFP features across the conditions while controlling the family-wise error rate at approximately α=0.05 using corrected harmonic mean p-values (HMP) (Wilson, 2018). This method, although not currently a standard practice, can test for differences in two samples in a way that is relatively insensitive to combining large numbers of tests (1856 in this case), allowing us to test for differences between groups without first combining power features into a small number of hand-picked frequency bands. The harmonic mean p-value approach is also robust to dependencies between tests, for example, those comparing power features in nearby frequency bins, and maintains reasonable statistical power. More specifically, we perform independent two-sided *t*-tests to compare the within-mouse averages of each LFP feature (power and cross-power) across windows. We then combine the p-values and calculate a “headline” p-value associated with the null hypothesis that the LFP features are the same across condition using the corrected harmonic mean p-value, following the procedure described by Wilson (2018). In comparison to procedures such as Benjamini-Hochberg that control the false discovery rate, the HMP is more sensitive in detecting significant groups of hypotheses at the expense of being less sensitive in detecting significant individual hypotheses (Wilson et al 2019). For this reason, we visualize the individual features with statistically significant differences between conditions by plotting only the mean differences in features that reach unadjusted significance at α=0.01, without regard to any HMP

### Experimental Design and Statistical Analysis

Data was plotted and analyzed in Graph Pad Prism 9.1 and custom Python scripts as described above. Sleep architecture comparisons for sleep medication were analyzed using RMANOVA followed by Holm-Šídák’s multiple comparison test and chronic light stress was analyzed using Wilcoxon matched-pairs signed rank test.

### Data and Code Availability

All data used for the analyses will be made available upon request to the corresponding author. All code used to preprocess LFPs, train factor models, make EMG-informed cluster-based sleep labels, and smooth labels is freely available online at https://github.com/carlson-lab/lpne (v0.1). Upon acceptance, this code will be input into the Duke University Research Data Repository, providing a permanent DOI and guaranteeing access for at least 30 years.

## Results

### Development and validation of brain-based sleep classifiers

To characterize oscillatory dynamics across the sleep-wake cycle, we implanted 9 C57BL/6J male mice with wires in 8 cortical and subcortical brain regions along with an EMG wire in the trapezius muscle. Mice had electrode wires targeting the cingulate cortex (CxCg), prelimbic cortex (CxPrL), infralimbic cortex (CxIL), nucleus accumbens (core and shell grouped together; NAc), amygdala (basolateral and central grouped together; Amy), medial dorsal thalamus (ThalMD), ventral hippocampus (HippV), and ventral tegmental area (VTA) (Figure 1A; Mague et al 2022). These regions were chosen given their well-established role in regulating emotional behavior under normal and pathological conditions (Nestler et al., 2002; Hultman et al., 2016; Hultman et al., 2018) and are routinely targeted with our microwire electrode arrays (Mague et al., 2022; Hughes et al., 2024). In order to determine the ground truth sleep-wake state label for each time window, we hand scored a random one-hour recording for wake, NREM and REM based on EMG power and CxPrL power based on known characteristics of these states (Keene and Duboue, 2018). We chose to bin the data into 2-second windows to reduce the number of time windows with multiple ground truth sleep labels while maintaining reliable spectral estimates. We established our accuracy benchmark based on inter-rater reliability from three raters on representative data used in this study where all datapoints were classified. The average Cohen’s kappa over 4 1-hour recordings and pairwise comparison of three independent raters was 0.82, range: 0.80-0.89; average agreement: 93.4% wake, 91.6%NREM, 63.1% REM. These results were in line with agreement typically reported between hand-scoring by different experts (Lee et al 2022).

**Figure 1.**
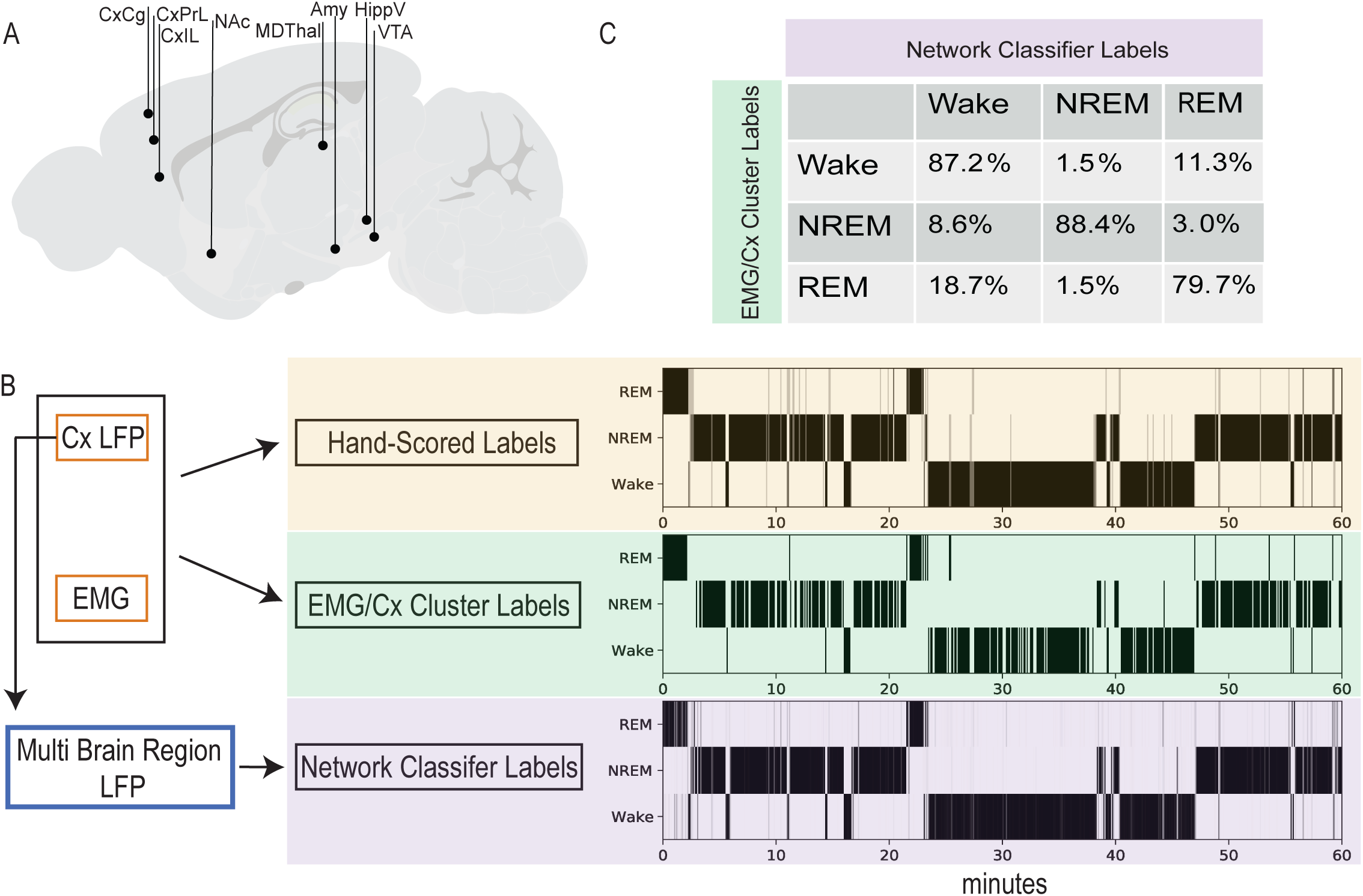
LFP-only network-based sleep labels recapitulate EMG-based labels. A. Recording sites from C57Bl6/J mice used for network training and recording. B. Diagram of input data used for three labeling methods and hypnodensity plots for a 1-hr segment showing classification of Wake, NREM, and REM based on hand-scored labels, EMG + Prelimbic cortex cluster-based labels, and network-based classifier labels. Intensity of shading indicates label confidence (e.g. agreement between human raters or network-model label probabilities). C. Confusion matrix of EMG + Prelimbic Cortex cluster-based labels in comparison to network-based classifier labels. Correct network labeling in comparison to EMG/Cx Cluster label percentages are seen in the diagonal for wake, NREM, REM respectively. Misclassification is reported in the off-diagonal. Misclassification is highest between wake and REM.

We then generated EMG-informed labels using traditional cortical LFP power and trapezius EMG power recorded over 24 hours per mouse by manual cluster identification, after validating these manual cluster-based labels with ground truth hand-scored labels (Figure 1B). Windows with features that did not clearly belong to a given cluster were left unclassified. EMG-informed labels were 95.5% in agreement for wake, 99.0% in agreement for NREM, and 91.6% in agreement for REM with the hand-scored labels from a randomly sampled hour of data from 7 mice. The human rater could not classify 12.6% of the data, whereas 17.3% of the data was unclassifiable using the EMG-informed method (n=4 male mice, 1 randomly sampled hour of recording data each). We proceeded with using EMG-informed labels for all training mice for further classifier training, since classification accuracy met the benchmark goal based on interrater reliability (Lee et al., 2022). First, we wanted to determine the ability of each emotion-regulating brain region to decode sleep on its own. We developed supervised autoencoder (SAE) classifiers to predict sleep-wake states. Each classifier was trained solely using spectral power from 1.9 to 54.7 Hz from one of the brain regions (CxCg, CxPrL, CxIL, NAc, Amy, ThalMD, HippV, and VTA. Classifiers for each of the brain regions examined could predict sleep-wake states above chance levels based on 33% balanced class accuracy being random chance (average wake accuracy: 73.6%, average NREM accuracy: 85.5%, average REM accuracy: 70.6%, p≤0.027 for each combination of sleep-wake state and region, one-sided Wilcoxon signed rank test comparing observed accuracy to 33%, n=9 based on nested cross-validation). While NREM accuracy was above 80% for all single-region decoders, the ability to distinguish between wake and REM was lower. Oscillations were most divergent across sleep-wake states in Amy, a previously reported region critical to gating REM sleep (Hasegawa et al., 2022), which gave the best single-region classification accuracy among sleep states (81% wake accuracy: 89% NREM accuracy, and 78% REM accuracy). In contrast, HippV oscillatory power was most similar across brain states and the produced the worst single-region classifier (58% wake accuracy: 84% NREM accuracy, and 56% REM accuracy). Thus, each emotion-associated region in isolation can decode wake, NREM, and REM states better than random chance with varying degrees of accuracy.

We hypothesized that the integration of information from all recorded sites would outperform single-region sleep state prediction, as we had previously observed (Grossman et al., 2024). To train a network classifier to predict sleep-wake states independently of EMG, we used the same ground truth labels used for the single-region classifiers and calculated power and synchrony features for frequencies up through 54.7 Hz from each of the recorded brain regions. We trained a supervised autoencoder (SAE) model to predict ground truth sleep labels based on LFP power and synchrony features (Talbot et al., 2023). Critically, this approach simultaneously reconstructs data and can predicts labels using a small number of latent dimensions. We found that the model had cross-validated accuracies of 87% for wake, 88% for NREM and 79.7% for REM, compared to the "ground truth" EMG-based approach (Figure 1C). Thus, the average recall of the classifier across all three sleep-wake states yielded an average balanced accuracy of 84.9%. This average balanced accuracy is higher than any of the single-region classifiers, and the network classifier performed significantly better than most single-region classifiers except for Amy (p=0.248) and NAc (p=0.082) (one-sided Wilcoxon rank sum tests, n=9).

We next determined the classifiers’ agreement with the EMG-informed labels in 5 new male hold-out mice. We found 86% ± 2% balanced accuracy (95% ± 2% agreement for wake, 71% ± 4% agreement for NREM, and 92% ± 4% agreement for REM; all values are mean ± SEM). In comparison to the mice used in cross-validation, there was no significant difference in balanced accuracy (p>0.999, two-tailed Mann-Whitney U test, n=5-9). We next wanted to test the discrimination of the multi-region classifier to the single region classifiers in hold-out subjects. We hypothesized that the single-region classifiers would not decode as well as the multi-region classifier, given our prior work (Mague et al., 2022). While the Amy and NAc classifiers had 71% and 75% balanced accuracy, respectively, these models (which had preformed similar to the network classifier during cross-validation) were significantly less accurate in holdout mice (p=0.0312 for both Amy and NAc compared to the network classifier, using one-sided Wilcoxon rank sum tests, n=5). Thus, the EMG-free network-based sleep-wake state classifier, and not the single-region classifiers, had similar or improved balanced accuracy in the holdout mice in comparison to the cross-validation analysis in the training mice. Due to this improved generalization, the network-based classifier was used for all following analyses.

Then we sought to test whether the network classifiers generalized to female mice. Female mice, in which both multi-site brain and EMG were recorded, exhibited high accuracy across the EMG-free network-based sleep-wake classifier (90 ± 4% accuracy for wake, 90 ± 3% for NREM, and 95 ± 4% for REM, 91.6% average balanced accuracy, n=4; all values are mean ± SEM) with statistically indistinguishable balanced accuracies from the cross-validated male accuracies (p=0.414, two-sided Mann-Whitney U test, n=4-9). Thus, the EMG-free network-based sleep-wake states classifier identified for sleep-wake states in male mice generalize to female mice.

### Power and synchrony profiles of distributed brain activity in wake, NREM, and REM

We then sought to describe the characteristic brain activity for each sleep-wake state across the monitored regions. To examine the brain activity during each state, we averaged power and synchrony across 24 hours for all timepoints identified for each state based on our network classifier. REM average power and synchrony features are more similar to wake than NREM features (0.94 vs. 0.84 cosine similarity), a difference that holds across all 9 mice used to train the classifier. NREM brain activity was dominantly composed of large delta (1-4Hz) activity without higher frequency activity (Figure 2B). Wake and REM also were composed of delta power across all regions and exhibited higher frequency activity (Figure 2A,C). We observed broad spectrum activity (2-52Hz) in HippV during wake and REM (Figure 2A,C).

**Figure 2.**
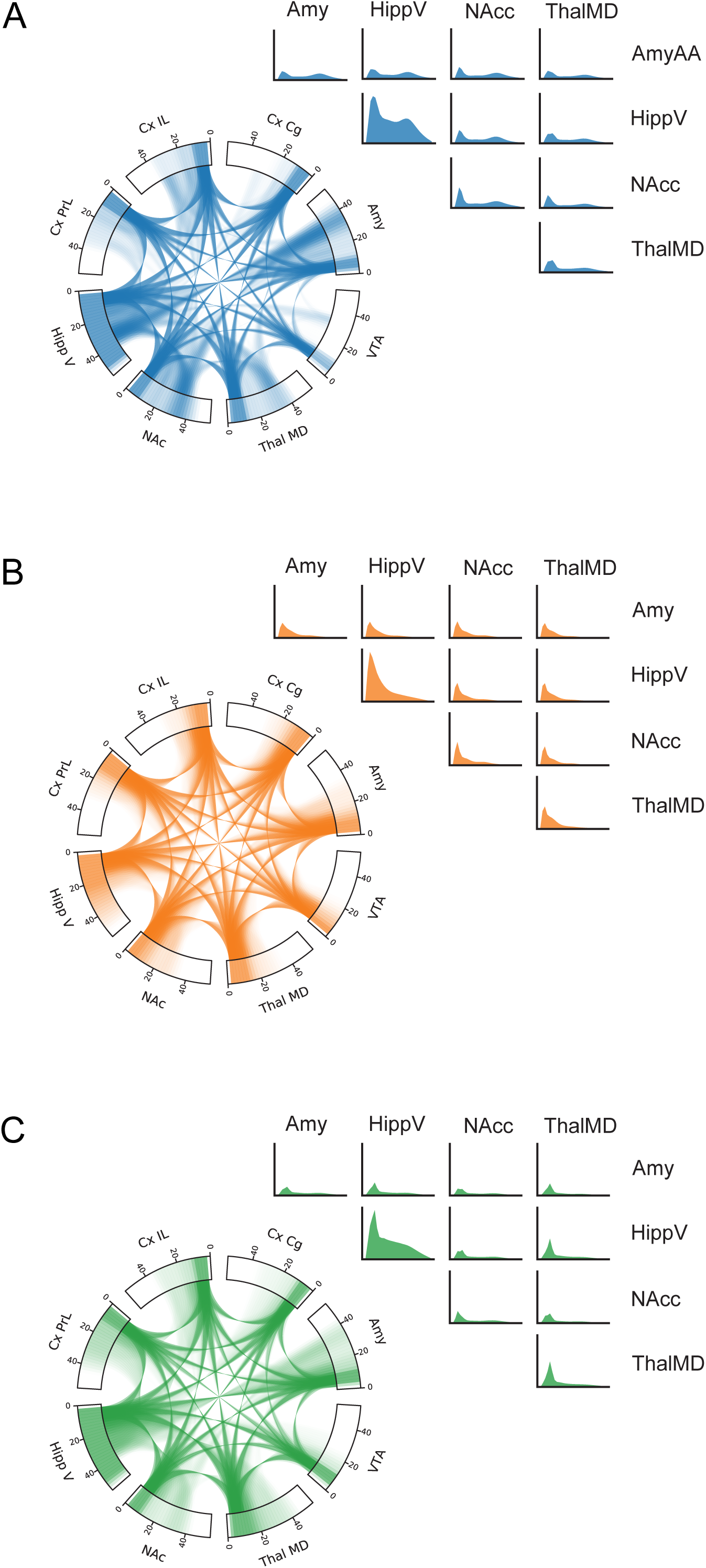
Average Brain Activity during wake, NREM, and REM. Chord plots showing power features from 0-55Hz along the outer rim for each brain region and connections between brain regions via connecting lines between brain regions in a frequency-dependent manner. Power (on the diagonal) and synchrony (on the off-diagonal) are plotted for each brain region for A. wake, B. NREM, and C. REM. Unthresholded average feature strengths plotted. N=9

Higher frequency (26-40Hz) power and synchrony was also observed in Amy, IL, PrL, NAc, and ThalMD during wake and REM, albeit more distinct during wake (Figure 2A,C). The average brain features from each state were consistent with the EEG-based brain oscillations that define sleep-wake states (Kohtoh et al., 2008).

To further disambiguate the sleep-wake states, we next examined the components of the supervised electome network for each of them. In contrast to the average brain activity in each state, the SAE models learn features that best discriminate the states from one another.

According to the model, wake was associated with increased power in Amy from 4-8Hz and 12- 55Hz as well as synchrony between Amy and HippV in the same ranges compared to the other two states (Figure 3A). NREM was associated with increased 10-55Hz activity in NAc and to a lesser extent to Amy as well as synchrony between NAc, CxPrL, and Amy (Figure 3B). REM was associated with increased 6-12Hz activity as well as 18-26Hz and 42-50Hz activity in ThalMD and HippV and to a lesser extent in VTA, Amy, and CxCg as well as synchrony amongst HippV, NAc, ThalMD, VTA, Amy, and Cg (Figure 3C). This finding extends widespread synchronous activity in sleep to subcortical regions including NAc, VTA, and Amy. Moreover, we establish the widespread synchronous activity profiles across brain regions for emotional regulation that are observed during distinct sleep states.

**Figure 3.**
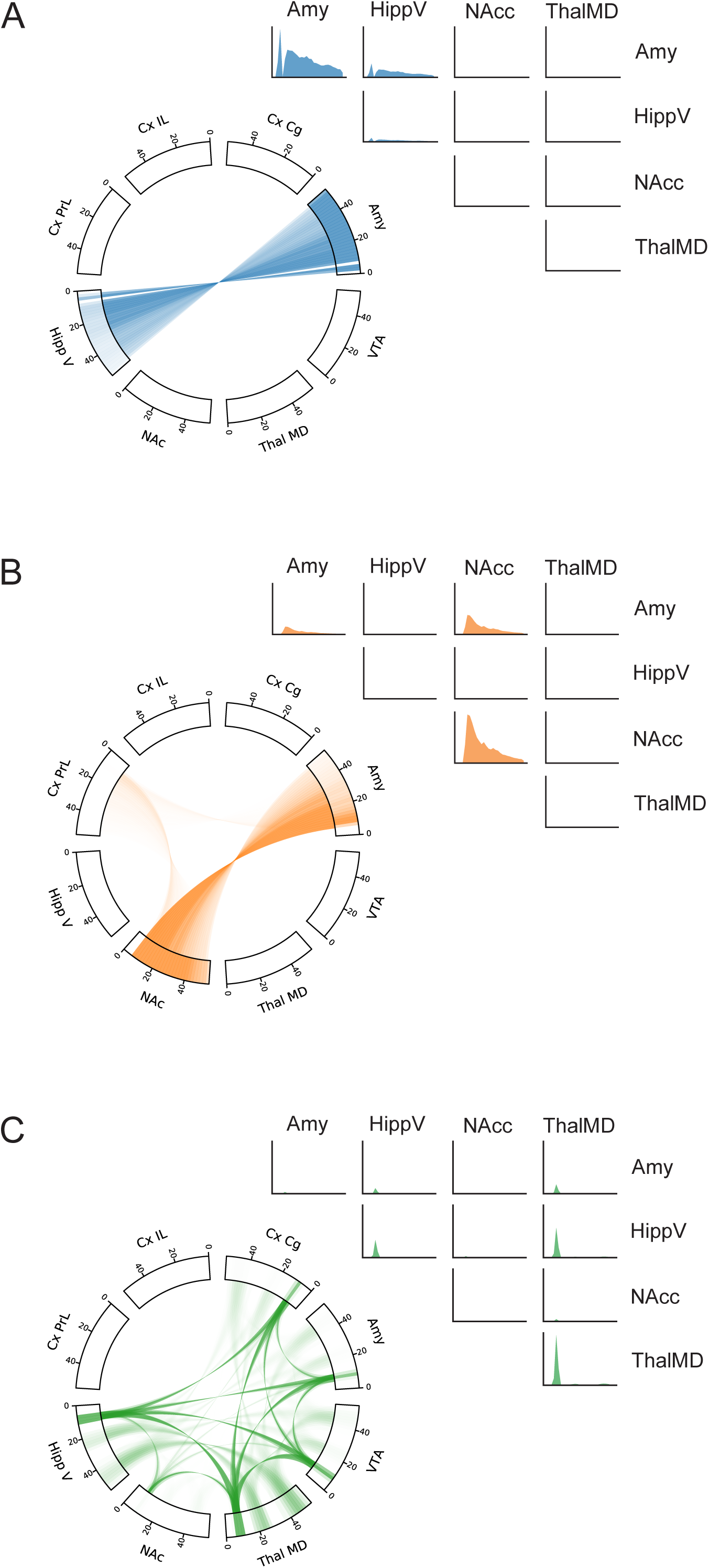
Electome Network Classifiers for sleep states. Chord plots show power features from 0-55Hz along the outer rim for each brain region and connections between brain regions via connecting lines between brain regions in a frequency dependent manner. Power (on the diagonal) and synchrony (on the off-diagonal) are plotted for each brain region for A. wake, B. NREM, and C. REM.

### Sex comparison shows no significant difference in brain dynamics during sleep

Even though the classifier generalized from male to female mice, it is still possible that male and female mice exhibit distinct oscillatory features during sleep that are separate from the features used by the classifier. We tested this hypothesis by examining the average power and synchrony features within the Wake, NREM, or REM epochs identified by the network classifier during 24-hour recordings of 13 male mice and 11 female mice. While NREM and REM had no significant differences between males and females (p=0.389 for NREM and p=0.403 for REM, corrected headline harmonic mean p-value across 1856 two-sided independent *t-*tests), wake brain activity pattern was significantly different (p=0.0497, corrected headline harmonic mean p- value across 1856 two-sided independent *t-*tests). Female mice on average exhibited increased synchrony between Amy and VTA from 38-52Hz while male mice on average had increased synchrony in the same projection from 30-34Hz during wake. Additionally, female mice had increased 10-12Hz synchrony between NAc and VTA and between Amy and VTA. Thus, NREM and REM brain activity is not statistically different between males and females whereas wake has significant sex differences in synchrony in sub-cortical brain structures.

### Trazodone and diazepam alter sleep architecture and brain dynamics across sleep-wake states

We next explored the widespread electrophysiological properties induced by two pharmacological agents commonly used to promote sleep—trazodone and diazepam. Drug concentrations were picked based on previous literature examining these drugs in mice (Kopp et al., 2003; de Oliveira et al., 2022). Trazodone decreases sleep latency and increases sleep duration with improved efficiency of sleep and increased time spent in NREM, in addition to serving as an antidepressant (Jaffer et al., 2017). The anxiolytic diazepam decreases sleep latency, increases sleep duration, and paradoxically also decreases the amount of REM (Holbrook et al., 2001). Thus, we examined the effects of these mood modulating agents on state-dependent brain dynamics. Here, we used a crossover design where trazodone, diazepam or a saline control was administered acutely.

To identify acute effects of trazodone and diazepam on sleep architecture, we first quantified sleep-wake states for the 4 hours after drug administration using the network-based classifiers. Indeed mice had significantly, or trends toward significantly, decreased NREM time (F_1.674,6.696_=12.77 RMANOVA P=0.0061; trazodone p=0.052, diazepam p=0.011, Holm-Šídák’s multiple comparisons test) and increased REM time for both drugs (F_1.462,5.848_=6.385 RMANOVA P=0.0391; trazodone p=0.030, diazepam p=0.028, Holm-Šídák’s multiple comparisons test), with no significant change in wake time (F_1.547,6.189_=1.186 RMANOVA P=0.3495) (Figure 4A). There was a significant decrease in average NREM bout length with both drugs (F_1.150,4.599_=32.24 RMANOVA P=0.0028; trazodone p=0.009, diazepam p=0.004, Holm-Šídák’s multiple comparisons test). However, there was a wide variation in bout length for REM after both trazodone and diazepam (trazodone P<0.01, diazepam P<0.01, F test to compare variances) (Figure 4B).

**Figure 4.**
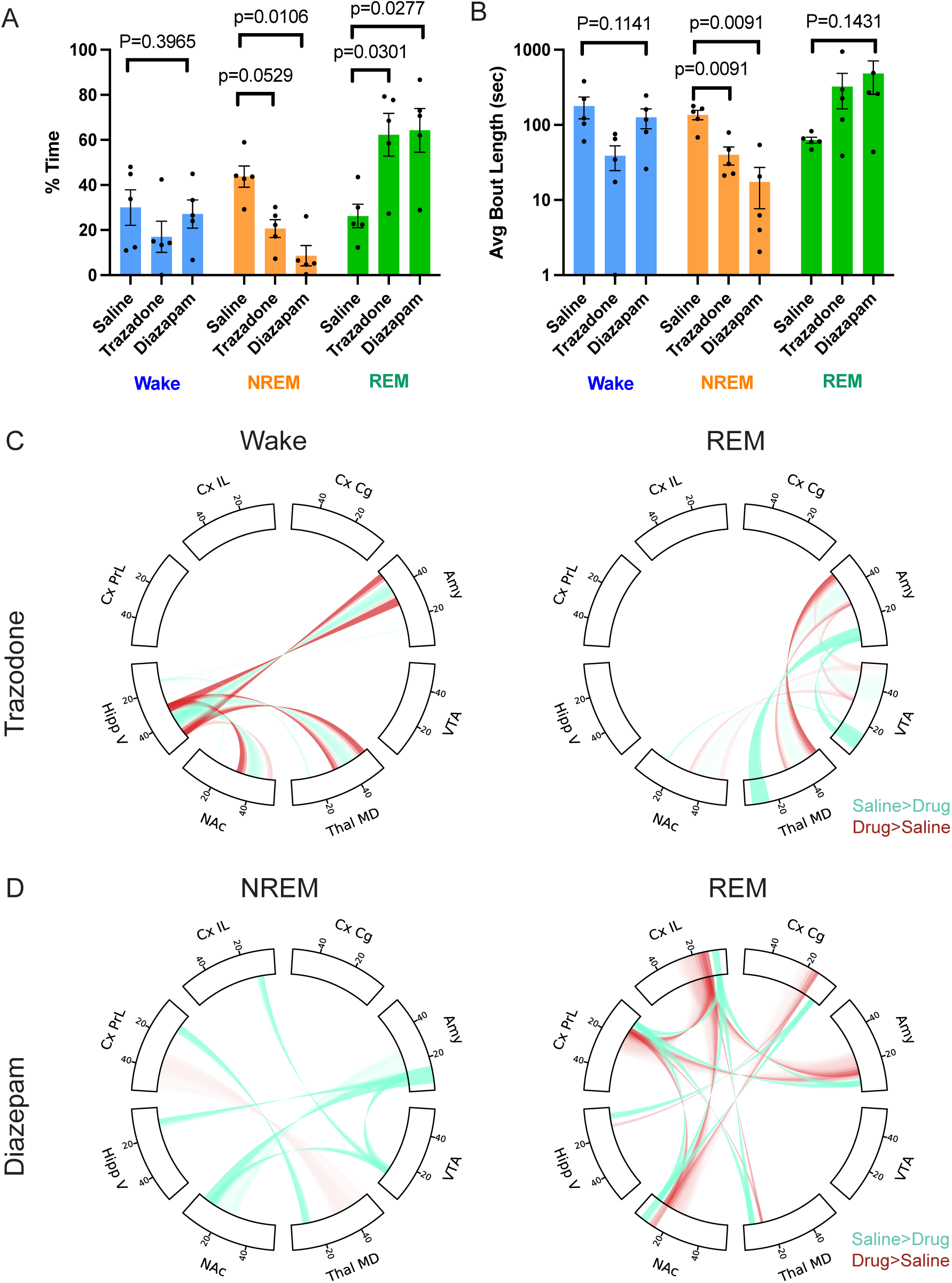
Brain-wide response to acute sleep medication. A. Trazodone and diazepam alter NREM (orange) and REM (green) but not wake (blue) overall time during the 4 hours after acute administration. (Wake: RMANOVA F_1.547,6.189_=1.186 P=0.3495; NREM: RMANOVA F_1.674,6.696_=12.77 P=0.0061; trazodone p=0.052, diazepam p=0.011, Holm-Šídák multiple comparisons test; REM: RMANOVA F_1.462,5.848_=6.385 P=0.0391; trazodone p=0.030, diazepam p=0.028, Holm-Šídák multiple comparisons test). B. NREM (orange) average bout length decreases in the 4 hours after acute administration of trazodone and diazepam but does not significantly alter bout length of wake (blue) and REM (green). (Wake: RMANOVA F_1.133,4.531_=3.792 P=0.1141; NREM: RMANOVA F_1.150,4.599_=32.24 P=0.0028; trazodone p=0.009, diazepam p=0.004, Holm-Šídák multiple comparisons test; REM: RMANOVA F_1.389,5.554_=2.877 P=0.1431). C,D. Significant power and cross power differences in the 4-hour period after acute trazadone (C) or diazepam (D) based on sleep network classifier state identification. Power and synchrony features that are increased in saline in comparison to drug are shown in teal, increased in drug in comparison to saline are shown in red. Threshold based on features that reach unadjusted significance at α=0.01. N=5.

We next sought to examine the brain dynamics across recording sites to examine acute effects of trazodone and diazepam during each sleep-wake state. Trazodone changed brain activity significantly during both wake and REM epochs (p=0.0417 and 0.0113 respectively, harmonic p-value) and brain activity changes trended toward significance for NREM (p=0.0662, harmonic p-value). During wake, trazodone caused increased synchrony between Amy/ThalMD/NAc and HippV in comparison to control around 30 and 50Hz but decreased around 40Hz (Figure 4C). During REM, trazodone increased synchrony around 30 and 50Hz and decreased synchrony around 40Hz between ThalMD and Amy, NAc and VTA, and VTA and Amy. Additionally, there was decreased power 6-16Hz in ThalMD and VTA as well as decreased synchrony in that same band between ThalMD and Amy (Figure 4C). Diazepam significantly altered brain activity during both NREM (p=0.0112, harmonic p-value) and REM (p=0.0020, harmonic p-value) but not wake (p=0.130, Holm-Šídák’s multiple comparisons test). Diazepam in comparison to control decreased 8-16Hz synchrony in CxIL and VTA, CxPrL and ThalMD, HippV and Amy, NAc and Amy, and NAc and VTA and a broad frequency (24-56Hz) increase in synchrony between CxPrL and ThalMD after drug administration during NREM (Figure 4D). REM increased around 14Hz synchrony between CxIL and CxPrL, CxPrL/CxIL and NAc, CxPrL/CxIL and Amy, and CxCg and NAc. There was decreased 6-10Hz synchrony among the same projections. Additionally, there was increased power around 14Hz and decreased power in comparison to control around 6-10Hz in CxIL and NAc (Figure 4D). Thus, trazodone and diazepam increase sleep and elicit differences in brain dynamics during sleep-wake states relative to sleep brain activity observed with control administration.

### Chronic circadian disruption alters sleep architecture without changing sleep brain dynamics

Finally, we examined sleep in a chronic stress-induced depressive-like model generated by aberrant light exposure to test if an emotion perturbation paradigm would change sleep quantity or content (Figure 5A, based on LeGates et al. (2012). Behavioral assessment of mice exposed to T7 light conditions for 14 days has been reported to resemble a depressive-like phenotype with decreased sucrose preference and increased immobility time during forced swim test (LeGates et al., 2012).

**Figure 5.**
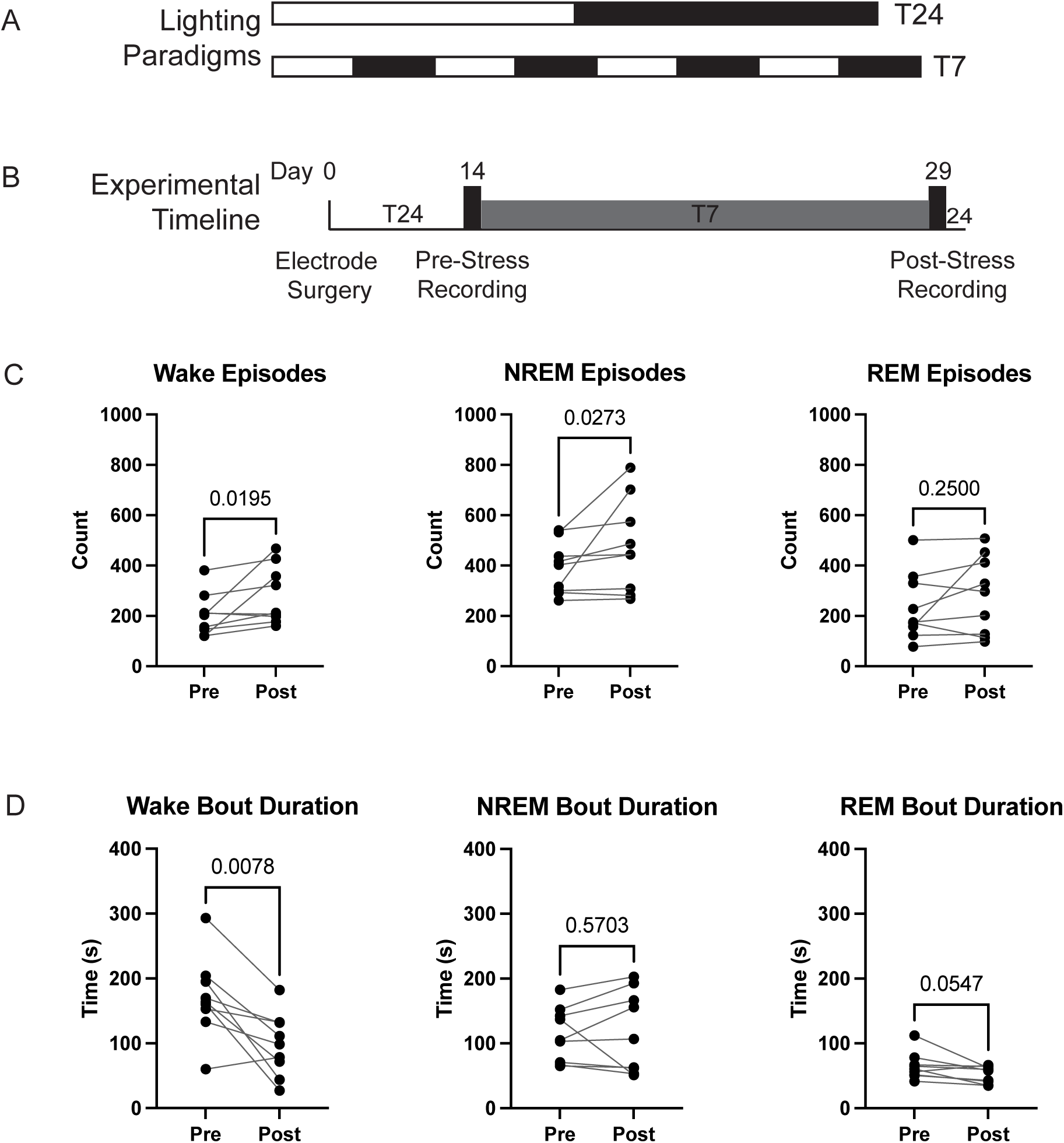
A. Scheme of light stress protocol. B. Timeline of experimental recordings. C. Increase in the number of episodes of wake and NREM (but not REM) after 14 days of T7 light stress. D. Significant decrease bout length of wake and trending to significant decrease in REM (but not NREM) bout duration following 14 days of T7 light stress. One-tailed Wilcoxon matched-pairs signed rank test. N=9.

We first examined the quantity of sleep by determining the overall percentage of time spent in each sleep-wake state over the course of 24 hours prior to stress and after 2 weeks of light stress (Figure 5B). Similar to prior reports, we observed no change to overall amounts of wake, NREM, or REM over the course of 24 hours (LeGates et al., 2012). However, after stress exposure, mice exhibited increased sleep fragmentation as indicated by increased number of episodes of wake and NREM (wake p=0.0195, NREM p=0.0273, Wilcoxon matched-pairs signed rank test; Figure 5C) and decreased bout length for wake (wake p=0.0078, Wilcoxon matched-pairs signed rank test; Figure 5D). Mice also tended to show a decrease in the bout length for REM, though these differences did not reach statistical significance (REM p=0.0547, Wilcoxon matched-pairs signed rank test; Figure 5D). Then we examined brain dynamics during each sleep/wake state by comparing power and synchrony across all recorded brain regions for each sleep classifier-based label. Averaging over the 24-hour pre-stress and post-stress periods, we observed no significant differences in power or synchrony across the brain (wake p=0.204, NREM p=0.261, REM p=0.361, corrected headline harmonic mean p-value across 1856 two-sided independent *t-*tests). Thus, chronic aberrant light cycle disruption induced sleep fragmentation abnormalities in mice with no difference in average brain dynamics of emotion-regulating regions during sleep states.

## Discussion

Here, we used a machine learning-based electome classifier to identify patterns of power and synchrony across emotion regulation brain region that distinguish wake, NREM, and REM states in mice, independent of electromyographic recordings. Oscillatory power in any one region, originally chosen due to their implication in emotional regulation or generation, could independently predict sleep-wake state at a level significantly higher than chance. We then developed classifiers using power and synchrony from all recorded brain regions and identified an electome model composed of a supervised component for each sleep-wake state, indicating that there are unique, stable patterns of brain activity across the observed regions during each sleep-wake state. This multi-region classifier performed significantly better than all single region classifiers and was conserved between sexes. We then examined brain dynamic perturbations by common psychotropic drugs known to modulate mood and sleep in humans and rodents. We found prominent differences between brain dynamics induced by both drug classes across sleep-wake states. Finally, we used the network classifier to identify disrupted sleep architecture in a chronic circadian stress-induced model of depression-like behavioral dysfunction. We found that chronic circadian stress increases sleep fragmentation, but the brain dynamics of each sleep state are indistinguishable in the pre- and post-stressed conditions.

A key methodological advance from our study is a novel strategy for sleep-wake classification from multi-region electrophysiological recordings. Sleep-wake classification using hand-scoring by trained experts, which remains common in the field, not only requires reliable EMG recordings, but is also time- and labor-intensive. Here we developed accurate classification of sleep-wake states utilizing EMG-free network classifiers from LFP recordings of several emotional regulatory brain regions. Mouse models of psychiatric disease are particularly susceptible to conditions that are incompatible with stable, long-term EMG recordings.

Longitudinal studies examining sleep require EMG for the duration of the study; however, the average duration of a quality EMG signal was 1-month post implantation, making any aging study or long-term longitudinal study impossible (such as in chronic stress paradigms e.g., Antoniuk et al. (2019)). Additionally, many psychiatric disease models are incompatible with quality EMG for various reasons including increased grooming (e.g. autism and obsessive-compulsive disorder, (Welch et al., 2007; Wang et al., 2016), physical exertion (aggression e.g., Golden et al. (2019), and excessive handling (restraint stress e.g., Mao et al. (2022) or social defeat e.g., Berton et al. (2006)). Similar to their human counterparts, traditional methods of sleep-wake state identification are not feasible in subjects that are unable to keep a muscle probe attached, which led our group and others to try to develop EMG-free or limited EMG classification approaches (Estrada et al., 2006; Ulrich et al., 2015; Bastianini et al., 2017; Kloefkorn et al., 2020; Zhang et al., 2022). However, these methods require time-consuming manual annotation of alternative signals such as whole-body plethysmography (Bastianini et al., 2017) or electrical field sensors (Kloefkorn et al., 2020) or are limited to cortical signals (Zhang et al., 2022). The development of non-EMG based classifiers of sleep-wake states allowed for tractable, long-term determination of sleep and wake in rodent models of emotional perturbation, opening the ability to examine mechanisms of sleep alterations without tedious manual annotations. Of the single-region models we developed, the Amy model was able to decode sleep-wake states in hold-out subjects, albeit not as well as the multi-region model. This suggests that Amy is a key brain region in sleep state identification in comparison to the other regions monitored in this study. This adds to the growing literature showing the importance of Amy in sleep where recent evidence showed that amygdala-in particular the basolateral amygdala- initiated the REM state (Hasegawa et al., 2022).

Secondly, our study continued to shed light on the interplay between sleep encoding in brain regions associated with emotion regulation. The sequence in which mental health conditions and sleep disruption emerge is unclear, but there is evidence to support that normalizing sleep can improve emotion regulation either before or after the emergence of a psychiatric condition (Goldstein and Walker, 2014; Jagannath et al., 2017). By better understanding the relationship between sleep and emotional regulation, we could generate better treatments to promote overall health. It will be important to conduct longitudinal studies over the course of disease progression to disentangle the relationship between sleep disruption and mental health symptom severity. Additionally, several classes of compounds commonly used to treat emotional disturbance are known to have sleep-promoting effects. The neural dynamics underlying these effects have not been previously examined. However, here we show that trazodone and diazepam increased the likelihood of brain activity being classified as REM after acute administration, and that the average brain dynamics classified during REM are altered differently by the two compounds. It is unclear if this altered REM brain state was as beneficial as unmedicated REM-like sleep for an organism. By developing drugs that promote sleep with normal sleep brain dynamics, we could disentangle sleep benefits from other effects of the medications.

Our network classifiers allowed us to investigate the sleep architecture in a mouse model of psychiatric disease. Indeed, we detected increased sleep fragmentation in a model of chronic stress induced behavioral adaptation generated by circadian disruption. Consistent with previous reports, the overall sleep quantity is unaffected (Altimus et al., 2008; LeGates et al., 2012).

However, further analysis revealed the fragmentation phenotype, in which state bout lengths were decreased. Whether the observed increase in sleep fragmentation represents a stress-related effect or a compensatory mechanism is an intriguing hypothesis to explore in future experiments. Additionally, since the brain dynamics during sleep were indistinguishable before and after the depression-like manipulation, normal sleep-inducing treatments may be beneficial at ameliorating sleep differences in those with depression. Interestingly, sleep reactivity, or the degree to which an individual’s sleep deteriorates when stressed, is correlated to the individual’s propensity for developing both pathological sleep disorders as well as other psychological disorders (Kalmbach et al., 2018) which highlights the translational value of studying mechanisms of stress-induced sleep disruption in mice.

Although average brain dynamics were unaltered during sleep-wake states in the model of chronic stress, other brain dynamic changes may exist. Namely, sleep-state transitions are disrupted in depression (Tsuno et al., 2005; Franzen and Buysse, 2008; Riemann et al., 2020). Indeed, we observed increased number of transitions post-chronic stress (Figure 5C, D). Now that we can rapidly classify the sleep-wake states, we could dissect the neural features that underly state transitions by examining the timepoints immediately preceding state change.

Identifying the processes that regulate the transitions among sleep states could reveal new avenues for intervention for sleep transition disruptions.

## Conclusions

In conclusion, we sought to determine the utility of emotion-associated regions for encoding sleep as well as the impacts of psychopharmacological and stress manipulations on their dynamics during sleep states. We developed an accurate, high throughput sleep-wake state classifier for brain-wide recordings in mice that can be applied across sexes and to a disease model. We validated the classifier and examined brain dynamics of emotion-associated regions during sleep states to compare female and male sex differences, emotional regulation medications, and stress-induced behavioral dysregulation. Sleep is a brain-wide phenomenon that impacts regions highly implicated in emotional disruption. Thus, the encoders described here allow for further dissection of the mechanisms comodulating sleep and emotional health.

## Acknowledgements

This work was supported by NIH R01MH120158, R01MH120158-S1 and Hope for Depression Research Foundation to KD, the American Academy of Child and Adolescent Psychiatry Pilot Research Award to ALB; and Hartwell Foundation Post-Doctoral Fellowship to KKW. We would like to thank Dr. Samer Hattar for consulting for the T7 Assay setup and helpful manuscript comments, Athy Robinson and Arielle Ramey for administrative support, Noah Lanier and Kat Hoefner for scientific discussions, and Dr. Jean Mary Zarate for helpful comments on the manuscript.

## Detailed Author Attributions

Kathryn K. Walder-Christensen – Experimental conceptualization and design for network training, female comparison, and sleep drugs with JG and ALB. Electrode construction for training EMG mice, female test mice, and sleep drug mice with ALB and SV. Implantation surgery on all training EMG mice, female test mice, and sleep drug mice. Data collection for training mice and female comparison mice with ALB and sleep drug mice with HS. Generated manual labels for hand label accuracy for 4 mice with ALB and SV. Performed sleep architecture analysis for sleep drug and T7 comparisons with RS. Performed histology for electrode track validation for all datasets with HS. Writing (initial draft and subsequent revisions) with ALB and JG. Funding acquisition (Hartwell).

Jack Goffinet– Developed all ML and GUIs for analysis (including manual sleep annotation GUI, feature generation GUI, model training, and model plotting). Designed and performed data analysis for sleep networks model tuning and manipulation comparisons (drug and T7) with KWC and RS. Writing (initial draft and subsequent revisions) with ALB and KWC.

Alexandra L. Bey– Conceived initial project with KD. Experiment conceptualization and design for network training, female comparison, and sleep drugs with KWC and JG. Data collection and preprocessing for original training mice & female comparison mice with KWC. Developed coding scheme for handscored labels in consultation with SK, handscored segments for 7 training mice, trained individuals who performed manual annotation of sleep wake states, electrode building for training mice and female mice with KWC, conducted sleep architecture analysis for training mice, funding acquisition (R01 supplement and AACAP pilot award), writing (initial draft and subsequent revisions) with KWC and JG.

Reah Syed- Processed data for T7 data analysis & sleep drug analysis with KWC, performed histology on training set mice with KWC.

Jacob Benton– Coordinated T7 data acquisition with SM and EA, built recording chambers and conducted T7 recordings.

Stephen D. Mague – Conducted T7 recordings with JB and EA and performed raw data curation. Performed surgery on sleep drug mice with KWC.

Elise Adamson- Conducted recordings on training mice with KWC and ALB and T7 recordings with JB and SM.

Sophia Vera – Electrode construction for sleep drug mice with KWC. Generated manual labels for hand label accuracy for 4 mice with ALB and KWC.

Hannah Soliman- Data acquisition for sleep drug experiment with KWC. Performed histology for electrode localization for all datasets with KWC and RS.

Sujay Kansagra- reviewed semi-automated labels for accuracy, gave feedback on modifications to make to semi-automated parameters, reviewed initial hand scored labels with ALB, paper editing.

David Carlson – Supervision of data analysis, funding, paper editing

Kafui Dzirasa – Conceived initial project with ALB, performed T7 mouse implantation surgeries, supervision of all project aspects. Designed and secured funding and resources, paper editing

